# CB1R blockade unmasks TRPV1-mediated contextual fear generalization in female, but not male rats

**DOI:** 10.1101/2023.04.12.536625

**Authors:** Kylie A Huckleberry, Roberto Calitri, Anna J. Li, Mackenna Mejdell, Ashna Singh, Vasvi Bhutani, Mikaela A Laine, Andrei S. Nastase, Maria Morena, Matthew N Hill, Rebecca M Shansky

## Abstract

Increasing evidence suggests that the neurobiological processes that govern learning and memory can be different in males and females, and here we asked specifically whether the endocannabinoid (eCB) system could modulate Pavlovian fear conditioning in a sex-dependent manner. Systemic (i.p.) injection of CB1R antagonist AM251 in adult male and female Sprague Dawley rats prior to auditory cued fear conditioning produced a female-specific increase in freezing that persisted across extinction and extinction retrieval tests but was prevented by co-administration of TRPV1R antagonist Capsazepine. Notably, AM251 also produced robust freezing in a novel context prior to auditory cue presentation the day following drug administration, but not the day of, suggesting that CB1R blockade elicited contextual fear generalization in females. To identify a potential synaptic mechanism for these sex differences, we next used liquid chromatography/tandem mass spectrometry, Western Blot, and confocal-assisted immunofluorescence techniques to quantify anandamide (AEA), TRPV1R, and perisomatic CB1R expression, respectively, focusing on the ventral hippocampus (vHip). Fear conditioning elicited increased vHip AEA levels in females only, and in both sexes, CB1R expression around vHip efferents targeting the basolateral amygdala (BLA) was twice that at neighboring vHip neurons. Finally, quantification of the vHip-BLA projections themselves revealed that females have over twice the number of neurons in this pathway that males do. Together, our data support a model in which sexual dimorphism in vHip-BLA circuitry promotes a female-specific dependence on CB1Rs for context processing that is sensitive to TRPV1-mediated disruption when CB1Rs are blocked.

## Intro

Over the last decade, recognition that fundamental brain functions can be mechanistically sex dependent has grown [1–4]. Within the field of learning and memory, recent fear conditioning studies suggest that male and female rodents not only engage unique behavioral repertoires during aversive learning and extinction [5–8], but also recruit discrete neural circuits and ensembles [9–12]. Because the principles of Pavlovian conditioning are frequently applied to cognitive behavioral therapies for disorders like PTSD [13], a better understanding of the neurotransmitter systems and cell signaling mechanisms that modulate associative learning and memory in both sexes will be critical to improving personalized approaches to mental health care.

One promising but under-studied system is the endocannabinoid (eCB) system. The two major eCBs in the brain, anandamide (AEA) and 2-Arachidonoylglycerol (2-AG), primarily act in a retrograde manner through presynaptic CB1 receptors (CB1R) on inhibitory terminals [14,15]. However, AEA can also act as an agonist for the transient receptor potential subfamily V member 1 (TRPV1) [16,17], a cation channel that is mostly known for its role in pain perception [18]. In addition, TRPV1 can play a modulatory role in emotional behavior and learning and memory processes, often with effects opposite those of the CB1 receptor [19,20]. Despite well documented sex differences in eCB signaling and receptor expression [21,22], the potential for CB1/TRPV1 dynamics to contribute to sex-dependent mechanisms of associative learning is poorly understood.

Recent findings by our team [23] suggest that specifically in female rats, AEA actions at CB1Rs may work to counter purportedly negative effects of TRPV1 signaling on fear extinction processes. Here we ask whether this interaction also applies to acquisition and consolidation of conditioned fear itself, with a particular focus on the potential for generalization in context encoding. To investigate a possible synaptic basis for the sex-specific effects of eCB manipulations we observe, we examine fear conditioning-elicited AEA levels in males and females and look for sex differences in expression of both TRPV1 and CB1Rs in the ventral hippocampus (vHip), a known hub for integrating contextual and emotionally salient information [24–26]. We find that CA1 vHip projections to the basolateral amygdala (BLA) are unique not only in their level of perisomatic CB1R expression, but also in that the density of the circuit itself is sexually dimorphic. This work therefore adds to increasing evidence that eCBs can mediate learning and memory in a sex-dependent manner, identifying a putative circuit through which these effects may occur.

## Materials and Methods

### Subjects

Adult male (n=115) and female (n=139) Sprague Dawley rats were used in this study (see Figure captions for individual group n’s). All animals were pair housed with food and water access ad libitum. The vivarium maintained a 12h light/dark cycle (lights on 0700) and all testing was conducted between 0900 and 1600. All procedures were conducted in accordance with the National Institutes of Health Guide for the Care and Use of Laboratory Animals and were approved by the Northeastern University Institutional Animal Care and Use Committee as well as in accordance with the Canadian Council on Animal Care (CCAC) guidelines and were approved by the University of Calgary Animal Care Committee.

### Behavior testing

Fear conditioning and extinction procedures were conducted in Coulbourn Instruments test chambers as described in [5–7]. The conditioned stimulus (CS) was a 30s 80dB 4Hz tone, and the unconditioned stimulus was a 1s 0.65mA footshock. Experimental design is shown in Figure 1A and was identical to that of our recent work [23]. Briefly, the protocol consisted of 10min habituation sessions in each context, followed by fear conditioning (7 CS-US pairs in context A), extinction (20 CS presentations in context B), and extinction retrieval (5 CS presentations in context B). Each session was conducted on consecutive days at the same time of day for each cohort. Animals whose tissue was used for Mass Spec or Western Blot analysis went through fear conditioning or were placed in the chambers and exposed to the CS only. All sessions were executed by Noldus Ethovision software, and recorded by video cameras mounted to the ceiling of the chamber. Behavioral data were collected by our custom Python tool ScaredyRat [5] combined with experimenter hand scoring to correct instances of animals sleeping during late extinction, which automated systems can score as freezing. All experimenters were blind to the animals’ drug condition.

**Figure 1.**
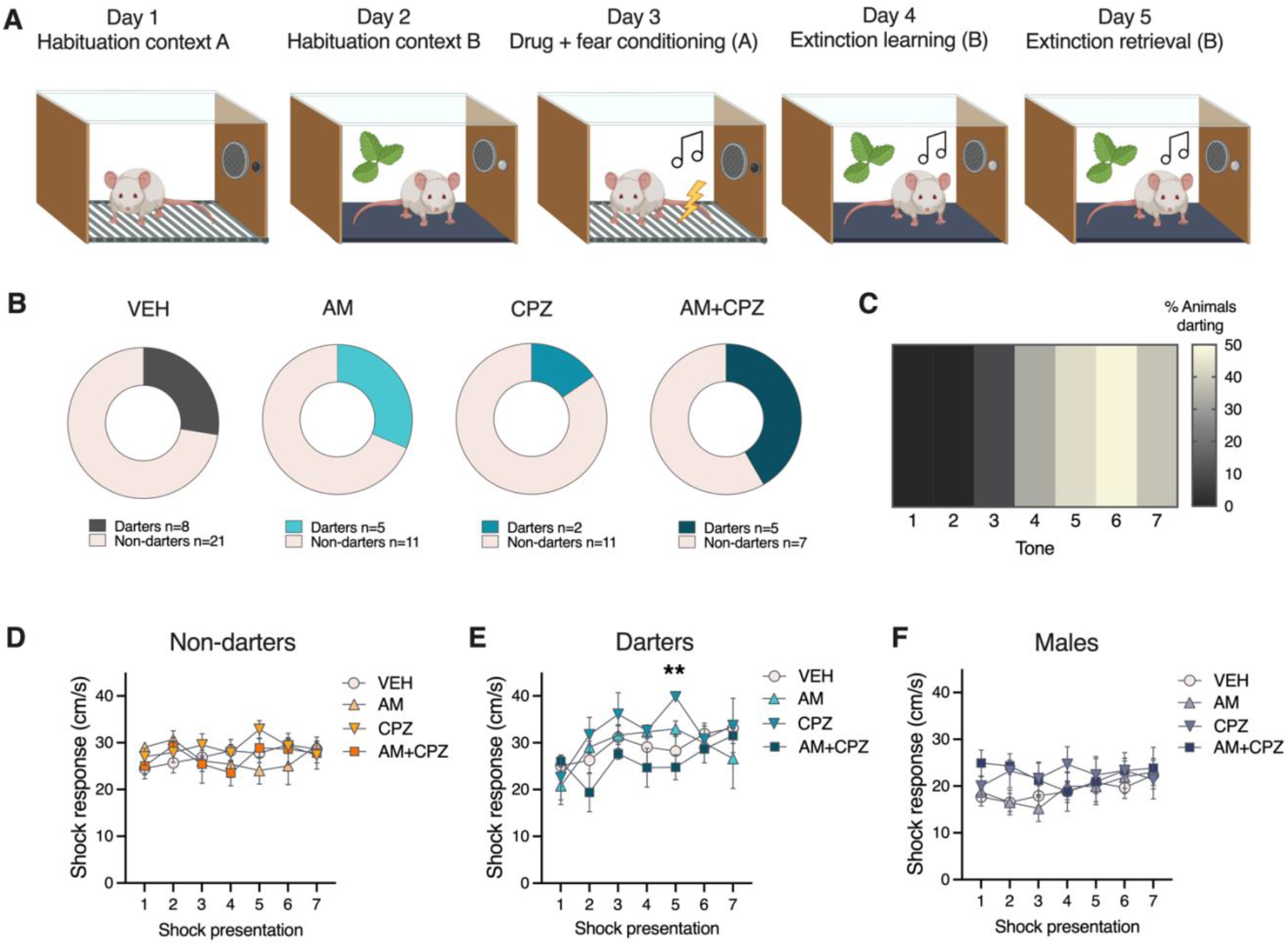
Experimental design, Darter proportions, and shock responsivity. A) Experimental design for behavior experiments consisted of 2 days of 10 min context habituation, drug administration before a 7 CS-US fear conditioning session, 20 CS extinction, and 5 CS extinction retrieval session. B) Proportion of animals that reached “Darter” criterion was not affected by drug administration. C) Heatmap showing conditioned darting prevalence by fear conditioning tone. D-F) Influence of drug treatment on shock responsivity (velocity at time of shock presentation) was observed only in Darters. **p<0.01 CPZ vs. AM+ CPZ, trial 5 only.

### Drug preparation

CB1 antagonist AM251 (1mg/kg; Tocris) and TRPV1 antagonist Capsazepine (5mg/kg; Cayman Chemicals) were prepared on the day of injection in vehicle solution (5% polyethylene glycol, 5% Tween-80, 90% saline), and were injected intraperitoneally at a volume of 1mg/ml 90 min before fear conditioning (Day 3). Doses were chosen to be identical to those used in [23].

### Euthanasia

Animals used for Mass Spec (MS) and Western Blot (WB) assays were euthanized via rapid decapitation 20 min after behavior testing ended. Animals in the naïve condition (WB only) never left their home cage and were euthanized at the same time of day. Immediately after euthanasia, brain regions of interest (for MS – medial prefrontal cortex [mPFC], amygdala, and ventral hippocampus [vHip]; for WB vHip only) were dissected, flash frozen, and stored at -80C.

### Liquid chromatography/Mass spectrometry

LC/MS was conducted as in [27]. Briefly, samples were homogenized in 2 mL of acetonitrile with 5 pmol d8-AEA (Cayman Chemical Company, Ann Arbor, Michigan, USA, #390050) in a borosilicate glass tube with a glass rod. Samples were sonicated, incubated overnight at -20 °C, and centrifuged at 1500×g. Supernatants containing lipids were isolated, evaporated with nitrogen gas, washed with acetonitrile and evaporated with nitrogen gas again. Final reconstitution for liquid chromatography/tandem mass spectrometry was in 200 μL of acetonitrile.

### TRPV1 Western Blot

vHip samples underwent subcellular fractionation protocols as in [28]. Cellular and membrane samples (50µg/subject) were then subjected to SDS-PAGE and Western Blot protocols using TRPV1 primary antibody (ThermoFisher PA1-29421; 1:1000). Primary antibodies for Beta-actin (ThermoFisher PA1-183; 1:1000) and Na^+^/K^+^ ATPase (MilliporeSigma 05-382; 1:1000) and associated secondaries (Vector PI-1000 and PI-2000; 1:1000) were used as respective loading controls for cellular and membrane samples. Optical density of all bands was imaged and quantified using ChemiDoc XRS (Bio-Rad) and Image Lab software. Bands were normalized by calculating the ratio of TRPV1 to loading control optical density for each animal. All data reflect the average of samples run in duplicate.

### Stereotaxic surgery

Experimentally naïve male (n=13) and female (n=14) rats underwent aseptic stereotaxic surgery for injection of 150nl retrograde-transducing AAV carrying fluorophore mCherry (pAAV-hSyn-mCherry; Addgene; 114472-AAVrg)) into the BLA (AP -3.0, ML+/- 5.0, DV -8.5). Animals were allowed to recover for four weeks to allow for viral transport and expression.

### CB1 immunohistochemistry

Rats were anesthetized with CO2 and transcardially perfused (flow rate: 45 ml/min) with cold 0.9% saline followed by cold 4% paraformaldehyde (PFA). Brains were post-fixed up to 36 hours in 4% PFA at 4°C. Serial coronal sections (50 μm) were cut through the entire basolateral amygdala and hippocampus on a vibratome (Leica Microsystems) and stored in 0.1% sodium azide in high molarity PBS. Four vHip-containing sections per animal were selected: sections ranged from A/P -4.68 mm to -6.48 mm [29]: A/P = -4.68 mm; A/P = -5.28 mm; A/P = -5.88; A/P = - 6.48). Free-floating sections were washed three times for 5 minutes in PBST (0.1% Triton-X in low molarity PBS) at room temperature on a shaker. Sections were next blocked in 5% normal donkey serum (NDS) in PBS for 1 h at room temperature on a shaker. Sections were then incubated with primary antibody (mouse anti-CB1R 1:500, Synaptic Systems, 258 011, 289C1, Lot 1-4) in blocking solution on shaker overnight at 4°C. Sections were washed the next day 3x 5 minutes each in PBST at room temperature on the shaker and incubated with a secondary antibody (Donkey anti-mouse 647 1:250, Jackson Immuno, 715-605-150, Lot 153967; diluted in glycerol per instructions) in 2.5% NDS in PBST for 2 h at room temperature on the shaker. Sections were next washed 3 times for 5 minutes in PBS at room temperature on the shaker. All sections were then mounted out of PBS-B in a petri dish onto Superfrost plus microscope slides (Fisherbrand, 12-550-15) and coverslipped (Epredia, 102455) with Fluoromount-G mounting medium (SouthernBiotech, 0100-01). Slides were stored at 4°C until imaging.

### Quantification of vHip-BLA projections

Fluorescent images of vHip CA1 were taken (Keyence Microscope, BZ-X710) using a 4x objective (PlanFluor 0.13 NA, PhL; 3s exposure) with Zoom x1.0. Images were then run through a custom ImageJ Macro (Github: kah218) using Fiji (ImageJ 2.3.0/1.53p). An experimenter then traced the vHPC, and using watershed and particle analysis, the macro quantified the number of labeled cells in the identified region. The counts and area of the traced region were then run through a custom Python script (Github: kah218) to generate the density of labeling in each section for each animal. Totals for each animal were averaged across sections.

### Confocal imaging

All images for CB1R quantification were acquired using an Olympus FV1000 confocal microscope (Optical Analysis Corporation). Cells were imaged using a 60x oil lens (UPLSAPO), 1.35 NA, Zoom of 3.7, and 0.33 µm step size. Using a 512 × 512 raster, these settings resulted in a resolution of 0.111 µm X 0.111 µm X 0.33 µm per pixel. Z-stacks of individual cells were acquired with a Kalman filter. For mCherry-labeled cells, images were acquired using a 543 argon laser, whereas images of CB1R immunohistochemical staining (AlexaFluor 647), images were acquired using a 635 argon laser. Approximately 4 mCherry-labeled and 4 unlabeled cells were imaged per section. Images to set background fluorescence were captured from the ipsilateral cortex for each section.

### CB1R quantification

Images were processed by a custom ImageJ Macro (Github: kah218) using ImageJ 1.53p. In short, this macro captures a “donut” from each Z-stack, comprising 5µm of perisomatic space. CB1R fluorescence was quantified by binarized particle analysis. These data were then processed by a custom Python script to determine normalized CB1R density (raw density – background density) values for each cell. Values for each cell type (mCherry-labeled vs. unlabeled) were averaged within sections and then across sections.

### Statistical analysis

All analyses were conducted with Graphpad Prism 9 software. Statistically significant group differences were detected via t-test, 1- or 2-way ANOVAs with post-hoc tests when appropriate. Specific tests used for each experiment are noted within the **Results** text below.

## Results

In the interest of brevity, we mostly only report statistical details for significant results here. Full statistical analyses can be found in Table S1. The design for Experiment 1 is shown in Figure 1A. We first examined our cohorts for the expression of conditioned darting [5,7]. Consistent with previous reports, 29% of females met the criterion to be classified as Darters, while no males did (Fig 1B). The proportion of Darters in each treatment group did not differ significantly (chi-square=2.2, p=0.53), suggesting that the drugs administered did not affect the propensity to dart. Corroborating prior reports [5,7,8], darting was most prevalent during later trials (Fig 1C). We next separated females into Non-darters and Darters, and evaluated shock response (e.g. the velocity at which an animal moves at the time of shock delivery) across trials. 2-way ANOVAs revealed no effects of trial or drug in Non-darters and Males (Fig 1D&F). In contrast, we found main effects of both trial (F_4,78_=2.6, p=0.04) and drug (F_3,118_=2.7, p<0.05) in Darters (Fig 1E). This main effect of time recapitulates previously observed patterns of progressively increasing shock response in Darters [5]. Corrected Dunnett’s post-hoc tests did little to identify a specific source of the main drug effect, revealing a significant difference only between CPZ and AM+CPZ groups at Tone 5 (p<0.01).

Figure 2 shows CS freezing data for Non-darters, Darters, and Males in all experimental groups across fear conditioning, extinction, and extinction retention. In Non-darter females (Fig 2A), we found main effects of both trial (F_5,236_=56.4, p<0.0001) and treatment (F_3,47_=2.7, p=0.01) during fear conditioning.

**Figure 2.**
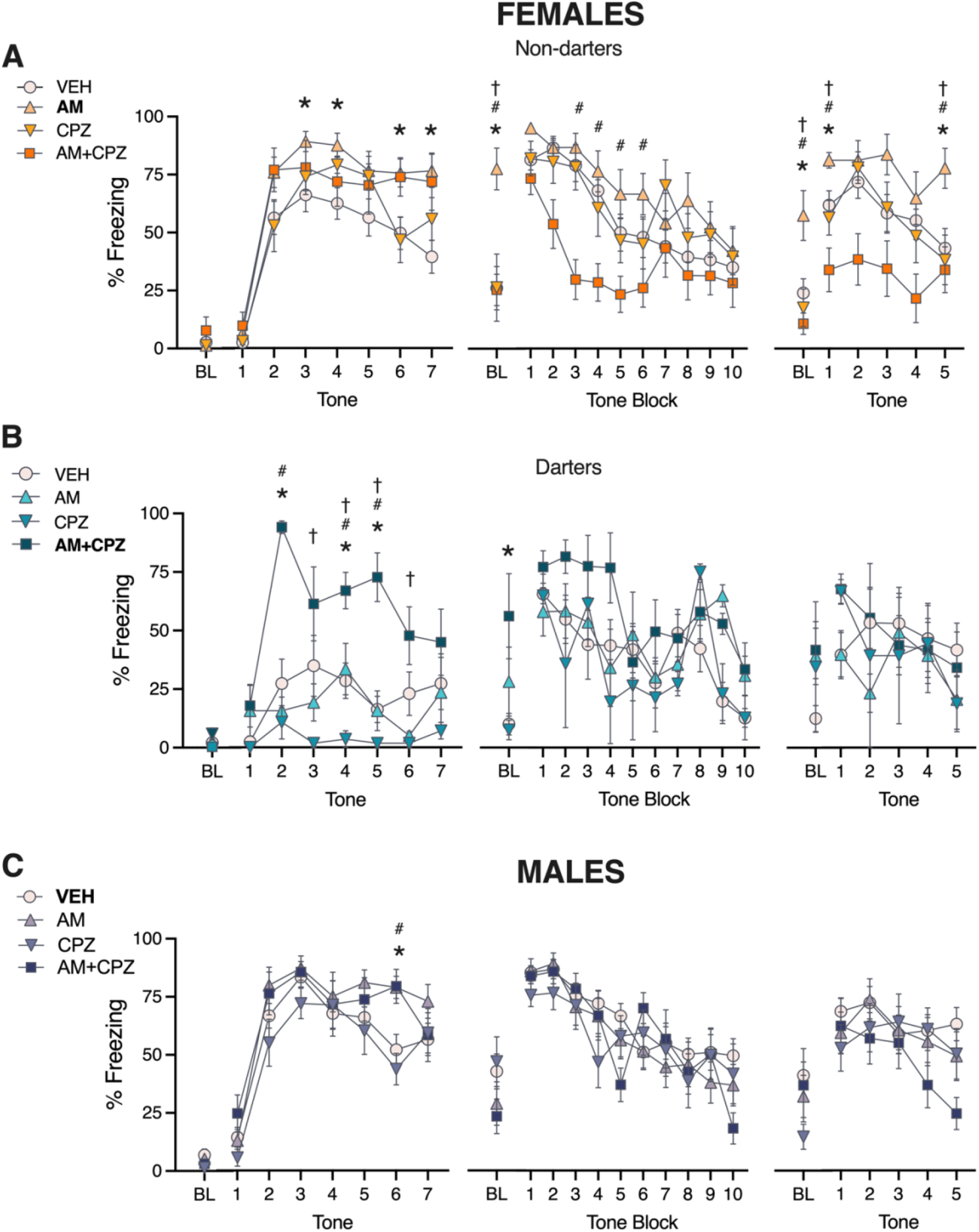
AM251 affects conditioned freezing in Non-darter females only. A-C) Conditioned freezing across Fear Conditioning, Extinction, and Extinction Retention in Female Non-darters, Female Darters, and Males (all Non-darters). Bolded treatment in each legend represents comparison of interest for post-hoc analyses. A) * AM vs. VEH; # AM vs. AM+CPZ; † AM vs. CPZ. B) * AM+CPZ vs. VEH; # AM+CPZ vs. AM; † AM+CPZ vs. CPZ. C) * VEH vs. AM; # VEH vs. AM+CPZ. For visual simplicity, single symbols are used for all p values less than 0.05. Detailed results are described in manuscript text and Table S1.

Dunnett’s post-hoc tests revealed that animals that received AM251 exhibited elevated freezing compared to vehicle during tones 3, 4, 6, and 7. No other group comparisons reached significance. On Day 2 (extinction), AM-treated females exhibited a robust freezing response during the baseline (BL) period compared to all other groups (1-way ANOVA: F_3,47_=7.4, p=0.0003; all post-hocs p<0.01), suggesting that AM treatment prior to fear conditioning resulted in contextual fear generalization the following day. Notably, this effect of AM251 was prevented by co-administration of TRPV1 antagonist CPZ. A 2-way ANOVA of freezing to extinction tones revealed main effects of both trial (F_9,423_=14, p<0.0001) and treatment (F_3,47_=4.02, p=0.01). Dunnett’s post-hoc tests revealed that AM+CPZ-treated animals exhibited an accelerated extinction learning curve (p≦0.01 vs. VEH and AM for trial blocks 3 & 4). Interestingly, these patterns persisted on Day 3 (extinction retention; main effect of treatment:). AM-treated animals again exhibited heightened freezing during the baseline period compared to all other groups (1-way ANOVA: F_3,47_=5.4, p=0.002; all post-hocs p<0.01). Additionally, AM-treated animals exhibited elevated freezing to tones across the session, while AM+CPZ animals exhibited suppressed freezing (p<0.05 AM vs. AM+CPZ all tones).

In Darters, a different pattern emerged (Fig 2B). While we again observed significant main effects of both trial (F_4,67_=2.7, p=0.03) and treatment (F_3,17_=11.4, p=0.0002) during fear conditioning, the latter effect was driven by AM+CPZ-treated animals, which exhibited a heightened freezing behavior uncharacteristic of Darters [5–7]. On Day 2, AM+CPZ animals exhibited elevated freezing during the baseline period (1-way ANOVA: F_3,17_=3.4, p=0.04; Dunnett’s post-hoc p<0.02 vs. VEH), but during CS presentation we found a main effect of trial only (F_4,66.5_=5.5, p=0.0008). No significant treatment effects were observed during extinction learning, nor during extinction retention.

In Males, we found significant main effects of trial (F_5,275_=41.5, p<0.0001) and treatment (F_3,54_=3, p=0.04) during fear conditioning. Dunnett’s post-hoc tests revealed a significant difference between VEH-treated animals and both AM and AM+CPZ-treated groups during tone 6 only (p<0.01 and 0.03, respectively). Significant main effects of time were found for both extinction (F_4,239_=18.6, p<0.0001) and extinction retention (F3,173=3.8, p=0.009), but no other significant treatment effects were observed.

Together, these data suggest that systemic manipulation of endocannabinoid signaling during fear conditioning induces effects that are not only sex-specific, but also behavioral phenotype-specific. CB1 blockade with AM produced long lasting fear generalization in a novel context in Non-darter females, but not Darters or Males. Non-darter females were also uniquely responsive to co-administration of CPZ, which reversed the AM-induced generalization and accelerated extinction learning and retention. In contrast, AM+CPZ produced elevated freezing in Darters and had little to no effect in males.

To begin to investigate a potential mechanism underlying the different responses to CB1R and TRPV1 manipulation in males and females, we first asked whether fear conditioning itself could induce sex-specific changes in AEA, which, unlike 2-AG, can bind to both receptors. Male and female rats underwent fear conditioning or CS exposure only in the conditioning context, followed by rapid decapitation and dissection of major brain regions known to be involved in fear conditioning and context processing, the medial prefrontal cortex (mPFC), amygdala, and ventral hippocampus (vHip; Figure 3A). We then performed liquid chromatography/tandem mass spectrometry to quantify tissue levels of AEA. In males, fear conditioning (CS + US) elicited an increase in mPFC AEA (t=2.6, p=0.02), but no conditioning-related changes were observed in the Amygdala or vHip (Figure 3B). In contrast, fear conditioning elicited an increase in AEA only in the vHip of females (t=2.4, p=0.02; Figure 3C).

**Figure 3.**
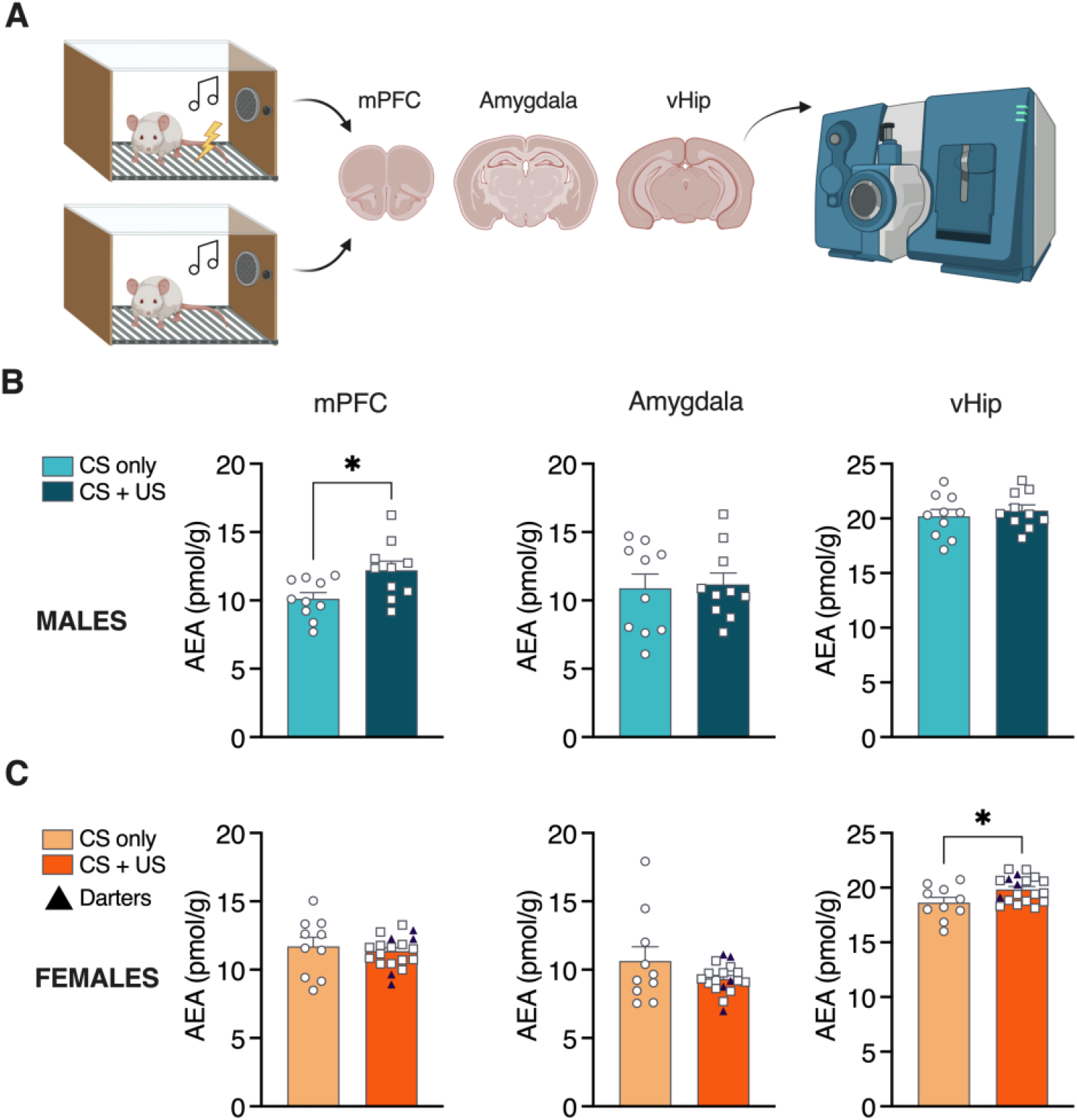
Fear conditioning induces sex- and location-specific increases in AEA. A) Experimental design for liquid chromatography/tandem mass spec quantification of AEA after fear conditioning. B) Fear conditioned males exhibited elevated AEA levels in mPFC. C) Fear conditioned females exhibited elevated AEA in vHip only. Darters were evenly distributed throughout fear conditioned samples for all brain regions assayed. *p<0.05 compared to CS only controls. Male n = 20 (10/10); Female n = 29 (10/19).

These data point to the vHip as a potential site of female-specific AEA signaling after fear conditioning. The vHip has been implicated in context fear generalization [30,31], and so to further investigate sex-specific synaptic mechanisms mediating our observed behavioral effects, we performed Western Blots in membrane fractionated vHip tissue, to allow detection of rapid insertion of TRPV1 receptors into cell membranes.

Figure 4A shows the design for this experiment. Animals were either fear conditioned as in the previous experiment (CS+US), exposed to the CS only, or remained in their home cages (naïve). Brains were extracted 20 min after the test ended, and vHip samples were dissected, fractionated, and run through standard Western Blot protocols for detection of TRPV1 receptor protein in both cytoplasm and membranes (Figure 4B). No main effects of group or sex were observed in either (Figure 4C-D). We also examined the ratio of cytoplasm:membrane TRPV1 as a potential measure of receptor mobility. We did not observe any significant group or sex effects (Figure 4E), suggesting that the effects we observed in Figure 2 may be due to sex differences in CB1 expression or mechanisms.

**Figure 4.**
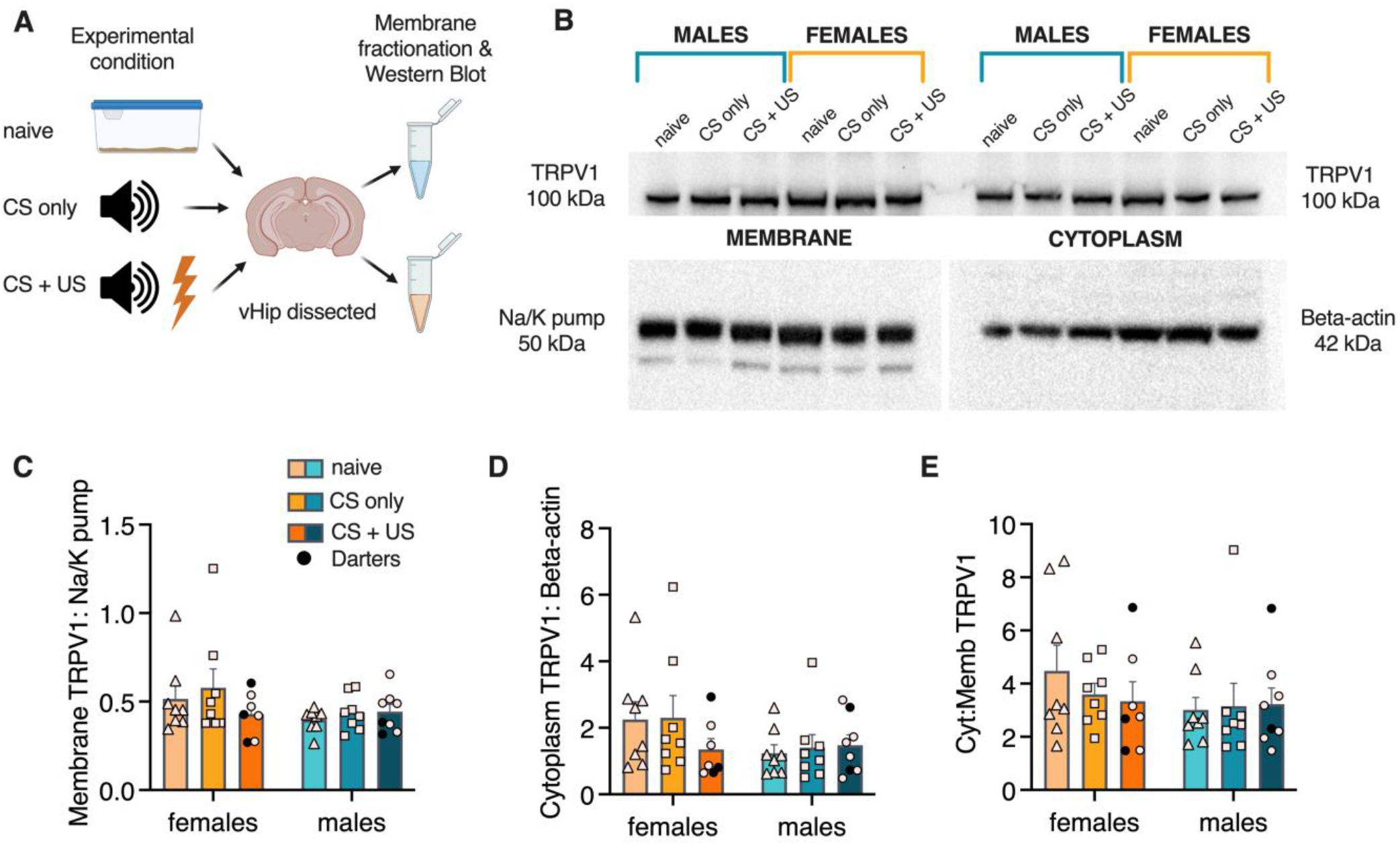
No sex differences in ventral hippocampal TRPV1 expression. A) Experimental design for membrane fractionated Western Blot for vHip TRPV1 expression. B) Representative blot showing lane organization and TRPV1 expression in membrane (left) and cytoplasm (right). Bottom blots show expression of control proteins for membrane and cytoplasm, respectively. C) Quantification of membrane TRPV1 expression, represented as proportion of Na/K pump expression. D) Quantification of cytoplasm TRPV1 expression, represented as proportion of Beta-actin expression. E) Ratio of cytoplasm to membrane TRPV1 expression. No significant main effects of sex or experimental condition or interactions were observed.

We next quantified perisomatic CB1R expression in vHip neurons of male and female rats, with the intent of examining the possibility that CB1R-expressing terminals might differ as a function of the vHip projection targets. The vHip-BLA pathway has been shown to be critical for context encoding in fear conditioning [32,33] and so we first injected a retrograde-transducing AAV-GFP into the basolateral amygdala (BLA; Figure 5A) and observed robust retrograde labeling in area CA1 (Fig 5B-C). We counter-stained vHip-containing sections for CB1R and observed perisomatic CB1R expression, as expected (Fig 5D). We next quantified perisomatic CB1R on both retro-labeled (BLA-projecting; Fig 5E) and unlabeled neurons by collecting Z-stacks of CB1R expression at selected cell bodies (Fig 5F) and, using custom Python scripts, determining the total level of fluorescence intensity within a standardized 3D “donut” around each soma (Fig 5G). In both sexes, vHip-BLA neurons contained significantly more perisomatic CB1R input than unlabeled cells (main effect of circuit, F_1,24_= 66.56, p<0.0001; Fig 5H).

**Figure 5.**
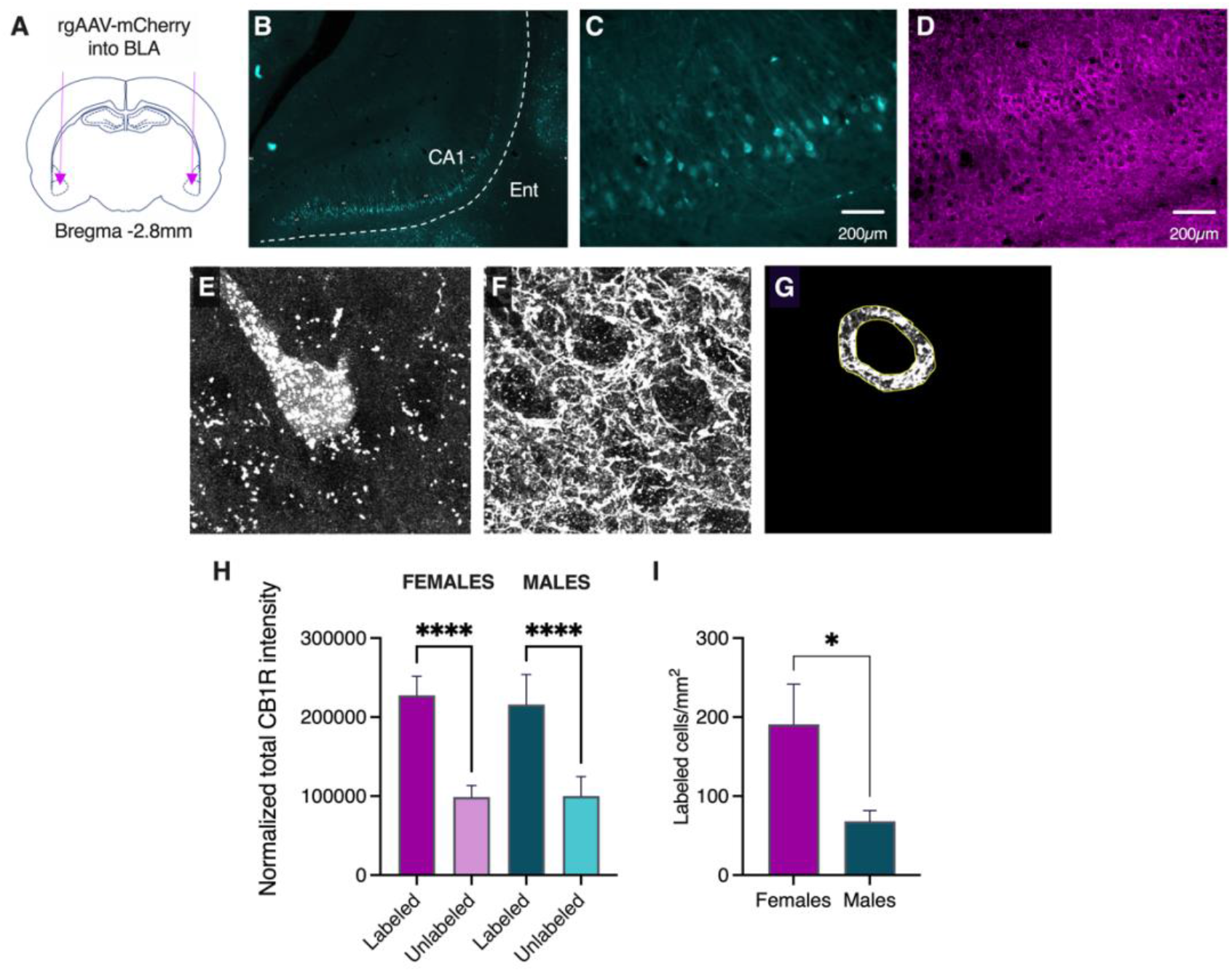
Perisomatic CB1R expression is twice as great at vHip-BLA neurons than at other vHip neurons. A) Surgical approach to retrogradely label vHip-BLA projections B) Representative micrograph showing viral expression in vHip CA1 C) Higher resolution micgrograph of retrogradely labeled vHip-BLA neurons D) Representative micrograph showing immunofluorescence for CB1R in vHip CA1 E) Representative confocal identification of retro-labeled vHip-BLA neuron. F) Retrograde confocal z-stack of perisomatic CB1R. G) Example of perisomatic “donut” captured by our custom Python script. H) Normalized total CB1R intensity in retro-labeled and unlabeled vHip neurons. I) vHip-BLA neurons were twice as abundant in females compared to males. ****p<0.0001; *p<0.05.

Finally, we quantified the number of vHip-BLA neurons in both sexes, finding that this pathway contained over twice as many cells in females as compared to males (Welch’s corrected t-test=2.32, p=0.03; Fig 5I).

Together, these data suggest that the vHip-BLA circuit is both sexually dimorphic and unique within the vHip in its CB1R innervation, pointing to a potential synaptic mechanism via which CB1R blockade could produce female-specific effects on learning and memory.

## Discussion

Here we investigated the impact of CB1R blockade on fear conditioning and extinction in male and female rats. CB1R antagonist AM251 elicited a long-lasting increase in freezing to both the CS and a novel context in females, but not males, an effect that was rescued (and then some) by co-administration of TRPV1R antagonist CPZ. These results suggest that under normal conditions and specifically in Non-darter females, CB1Rs maintain healthy associative learning and context processing by countering potentially negative effects of TRPV1 signaling, which are unmasked by CB1R blockade. Our findings parallel those of our prior work [23], which found that pre-extinction administration of AM251 also elicited a female-specific increase in freezing that persisted across days and was rescued by co-administration of CPZ. One notable difference between that study and the present one is that the AM251 effect in the former was only observed when the FAAH inhibitor URB597 was also administered, suggesting that elevated AEA levels are required for TRPV1 signaling to impact learning and memory processes. Indeed, we find here that fear conditioning on its own is sufficient to raise AEA levels in the female vHip, identifying a potential catalyst for the female-specific response to CB1R and TRPV1 antagonism. In both studies, the behavioral effects of both AM251 and AM251 + CPZ extended beyond the day of drug administration, suggesting that these manipulations may be facilitating stable changes in synaptic strength. Supporting this idea is evidence from other labs demonstrating long lasting effects of TRPV1 blockade on conditioned fear in male mice, which exhibited reduced freezing and resistance to reinstatement over a week after drug administration [34].

Intriguingly, these pharmacological effects were only observed in animals that were classified as Non-darters. As we have reported previously [5–8], Darters here exhibited discrete patterns of behavior both in response to the unconditioned (shock) and conditioned (tone) stimuli, providing further evidence that Darters represent an identifiable phenotype with unique defensive behavioral repertoires. Most striking in these animals was the response to combined AM251 and CPZ administration, which resulted in elevated freezing during fear conditioning and early extinction. That this drug effect was directionally opposite to what we observed in Non-darters points to the eCB system as a potential mechanistic basis for divergent behaviors in Darters vs. Non-darters. We did not observe a difference in fear conditioning-related AEA levels in Darters, suggesting that these differences may be related to CB1R and/or TRPV1 expression. Ongoing work in our lab is testing this hypothesis.

vHip CB1R quantification produced two key insights into the potential role for the vHip-BLA pathway in our findings. First, vHip-BLA projections were twice as abundant in females as in males. This is, to our knowledge, the first demonstration of sexual dimorphism in efferent density of an isolated hippocampal circuit. vHip-BLA projections have been shown to be critically involved in multiple forms of associative learning, including context fear conditioning as well as observational fear learning. Specifically, the neurons that comprise the vHip-BLA circuit undergo synaptic potentiation as a result of fear conditioning, and are then recruited during recall to reactivate memory engram ensembles in the BLA [33]. Kim & Cho (2020) demonstrated that vHip-BLA engagement is context-specific, and that discrete subsets of vHip-BLA projections encode different contexts. In this light, we speculate that in females, the increased density of vHip-BLA projections makes it more likely that under certain circumstances, neurons encoding the “wrong” context could be activated. Our data suggest that one such circumstance may be the loss of CB1R signaling, which here resulted in contextual fear generalization and elevated cue-induced freezing through unmasking of TRPV1 signaling via AEA activity.

Our findings are in line with previous work in mice that identified stronger vHip-BLA synaptic connectivity as a biomarker of stress-susceptible subpopulations compared to stress resilient mice [35]. The susceptible phenotype appeared to be CB1R-dependent and was specific to vHip-BLA connectivity, as no differences between stress-susceptible and -resilient mice were observed in the strength of other BLA afferents. With the caveat that only male animals were used, these data and our finding that CB1Rs targeting vHip-BLA neurons are twofold that of other vHip neurons point to the vHip-BLA circuit as uniquely positioned to respond to alterations in CB1R signaling in ways that affect threat responsivity. More specifically, we propose that the combined increase in perisomatic CB1R expression and greater density of vHip-BLA projection neurons in females results in dysregulation of this pathway when CB1Rs are blocked, resulting in poor contextual representation of the fear conditioning memory.

Another potential contributor to our sex-dependent observations may be the influence of estradiol (E2) on eCB signaling in the vHip. Huang & Woolley [36] demonstrated that estradiol can elicit a suppression of GABA release on CA1 neurons by stimulating post-synaptic mobilization of AEA (but not 2-AG). This CB1R-dependent effect occurred via membrane-bound E2-alpha receptors and was specific to females, despite evidence that hippocampal estradiol levels are comparable across the sexes [37]. Additionally, *in vitro* work has shown that E2 can rapidly sensitize TRPV1Rs, leading to potentiated neuronal excitation in the presence of capsaicin, a TRPV1R agonist [38]. Estradiol is therefore positioned to promote TRPV1 activity by both increasing levels of an endogenous ligand and sensitizing the receptor to stimulation. In the context of the current study, we propose that while E2-driven AEA in females may be beneficial to learning and memory processes when CB1Rs are available, it may in fact be harmful in cases when it favors TRPV1R signaling. Testing this hypothesis with pharmacological or genetic manipulations of estrogen receptors will be an important next step in our work.

In sum, we found that systemic CB1R blockade before cued fear conditioning produced elevated conditioned and contextual freezing in Non-darter female rats, but not Darters or males. This effect lasted across both extinction and extinction retention sessions but was reversed by co-administration of a TRPV1R antagonist that had no effect on its own. While no sex differences in TRPV1R expression were observed in vHip cytoplasm or membrane fractions, we found that perisomatic CB1R at vHip-BLA projections was twice that of other vHip cells, and that this pathway was twice as dense in females as in males. Together, these findings point to vHip-BLA circuitry as a novel mechanism through which eCBs may elicit sex-dependent effects on learning and memory. Our work opens the door for more direct inquiries into sex differences in eCB modulation of vHip-BLA function, with the potential to identify novel, sex-specific therapeutic targets.

## Acknowledgements

We are grateful for the assistance of Prabarna Ganguly, Jennifer Honeycutt, and Lauren Granata from the Brenhouse lab, who provided training, protocols, and reagent lists for the Western Blot study. Thanks are also due to Jordan Abettan for drug preparation, and Sean Trettle for his work on ScaredyRat.

## Author Contributions

Experimental design: RMS, MNH, and KAH; Behavior testing and data collection: AJL, MM, AS; Mass Spec: AN and MM; Western Blot: RC; Stereotaxic surgeries, CB1R immunostaining, and imaging: KAH; Coding and macro development; KAH and MAL; Cell counting: VB; Data analysis: RMS, KAH, and RC; Manuscript prep: RMS, MAL, MNH.

## References

1. Mogil JS. Qualitative sex differences in pain processing: emerging evidence of a biased literature. Nat Rev Neurosci. 2020;21:353–365.

2. Melcangi RC. Structural and molecular brain sexual differences: A tool to understand sex differences in health and disease. Neurosci Biobehav Rev. 2016;67:2–8.

3. Bangasser DA, Valentino RJ. Sex differences in stress-related psychiatric disorders: Neurobiological perspectives. Front Neuroendocrinol. 2014;35:303–319.

4. Fenton GE, Halliday DM, Mason R, Bredy TW, Stevenson CW. Sex differences in learned fear expression and extinction involve altered gamma oscillations in medial prefrontal cortex. Neurobiol Learn Mem. 2016;135:66–72.

5. Mitchell JR, Trettel SG, Li AJ, Wasielewski S, Huckleberry KA, Fanikos M, et al. Darting across space and time: parametric modulators of sex-biased conditioned fear responses. Learn Mem. 2022;29:171–180.

6. Laine MA, Mitchell JR, Rhyner J, Clark R, Kannan A, Keith J, et al. Sounding the Alarm: Sex Differences in Rat Ultrasonic Vocalizations during Pavlovian Fear Conditioning and Extinction. ENeuro. 2022;9:1–48.

7. Gruene TM, Flick K, Stefano A, Shea SD, Shansky RM. Sexually divergent expression of active and passive conditioned fear responses in rats. Elife. 2015;4.

8. Colom-Lapetina J, Li AJ, Pelegrina-Perez TC, Shansky RM. Behavioral diversity across classic rodent models is sex-dependent. Front Behav Neurosci. 2019;13.

9. Urien L, Bauer EP. Sex Differences in BNST and Amygdala Activation by Contextual, Cued, and Unpredictable Threats. ENeuro. 2022;9.

10. Tronson NC. Focus on females: a less biased approach for studying strategies and mechanisms of memory. Curr Opin Behav Sci. 2018;23:92–97.

11. Keiser AA, Turnbull LM, Darian MA, Feldman DE, Song I, Tronson NC. Sex Differences in Context Fear Generalization and Recruitment of Hippocampus and Amygdala during Retrieval. Neuropsychopharmacology. 2017;42:397–407.

12. du Plessis KC, Basu S, Rumbell TH, Lucas EK. Sex-Specific Neural Networks of Cued Threat Conditioning: A Pilot Study. Front Syst Neurosci. 2022;16.

13. VanElzakker MB, Kathryn Dahlgren M, Caroline Davis F, Dubois S, Shin LM. From Pavlov to PTSD: The extinction of conditioned fear in rodents, humans, and anxiety disorders. Neurobiol Learn Mem. 2014;113:3–18.

14. Kawamura Y, Fukaya M, Maejima T, Yoshida T, Miura E, Watanabe M, et al. The CB1 cannabinoid receptor is the major cannabinoid receptor at excitatory presynaptic sites in the hippocampus and cerebellum. J Neurosci. 2006;26:2991–3001.

15. Dow-Edwards D, Silva L. Endocannabinoids in brain plasticity: Cortical maturation, HPA axis function and behavior. Brain Res. 2017;1654:157–164.

16. Starowicz K, Cristino L, di Marzo V. TRPV1 receptors in the central nervous system: potential for previously unforeseen therapeutic applications. Curr Pharm Des. 2008;14:42–54.

17. Katona I, Freund TF. Multiple Functions of Endocannabinoid Signaling in the Brain. Annu Rev Neurosci. 2012;35:529–558.

18. Rosenbaum T, Simon SA. TRPV1 Receptors and Signal Transduction. CRC Press/Taylor &Francis; 2007.

19. Uliana DL, Hott SC, Lisboa SF, Resstel LBM. Dorsolateral periaqueductal gray matter CB1 and TRPV1 receptors exert opposite modulation on expression of contextual fear conditioning. Neuropharmacology. 2016;103:257–269.

20. Moreira FA, Aguiar DC, Terzian ALB, Guimarães FS, Wotjak CT. Cannabinoid type 1 receptors and transient receptor potential vanilloid type 1 channels in fear and anxiety—two sides of one coin? Neuroscience. 2012;204:186–192.

21. Ney LJ, Matthews A, Bruno R, Felmingham KL. Modulation of the endocannabinoid system by sex hormones: Implications for posttraumatic stress disorder. Neurosci Biobehav Rev. 2018;94:302–320.

22. Viveros M, Llorente R, Suarez J, Llorente-Berzal A, López-Gallardo M, Rodriguez de Fonseca F. The endocannabinoid system in critical neurodevelopmental periods: sex differences and neuropsychiatric implications. Journal of Psychopharmacology. 2012;26:164–176.

23. Morena M, Nastase AS, Santori A, Cravatt BF, Shansky RM, Hill MN. Sex-dependent effects of endocannabinoid modulation of conditioned fear extinction in rats. Br J Pharmacol. 2021;178.

24. Hobin JA, Ji J, Maren S. Ventral hippocampal muscimol disrupts context-specific fear memory retrieval after extinction in rats. Hippocampus. 2006;16:174–182.

25. Rudy JW, Matus-Amat P. The ventral hippocampus supports a memory representation of context and contextual fear conditioning: implications for a unitary function of the hippocampus. Behavioral Neuroscience. 2005;119:154–163.

26. Czerniawski J, Ree F, Chia C, Otto T. Dorsal versus ventral hippocampal contributions to trace and contextual conditioning: Differential effects of regionally selective nmda receptor antagonism on acquisition and expression. Hippocampus. 2011;22:1528–1539.

27. Vecchiarelli HA, Morena M, Lee TTY, Nastase AS, Aukema RJ, Leitl KD, et al. Sex and stressor modality influence acute stress-induced dynamic changes in corticolimbic endocannabinoid levels in adult Sprague Dawley rats. Neurobiol Stress. 2022;20.

28. Ganguly P, Holland FH, Brenhouse HC. Functional Uncoupling NMDAR NR2A Subunit from PSD-95 in the Prefrontal Cortex: Effects on Behavioral Dysfunction and Parvalbumin Loss after Early-Life Stress. Neuropsychopharmacology. 2015;40:2666.

29. Paxinos G, Watson C. The Rat Brain in Stereotaxic Coordinates. 7th ed. Academic Press; 2013.

30. Cullen PK, Gilman TL, Winiecki P, Riccio DC, Jasnow AM. Activity of the anterior cingulate cortex and ventral hippocampus underlie increases in contextual fear generalization. Neurobiol Learn Mem. 2015;124:19–27.

31. Ortiz S, Latsko MS, Fouty JL, Dutta S, Adkins JM, Jasnow AM. Anterior cingulate cortex and ventral hippocampal inputs to the basolateral amygdala selectively control generalized fear. 2019;12:329–348.

32. Terranova JI, Yokose J, Osanai H, Marks WD, Yamamoto J, Ogawa SK, et al. Hippocampal-amygdala memory circuits govern experience-dependent observational fear. Neuron. 2022;110:1416–1431.e13.

33. Kim W Bin, Cho JH. Encoding of contextual fear memory in hippocampal–amygdala circuit. Nat Commun. 2020;11.

34. Iglesias LP, Fernandes HB, de Miranda AS, Perez MM, Faccioli LH, Sorgi CA, et al. TRPV1 modulation of contextual fear memory depends on stimulus intensity and endocannabinoid signalling in the dorsal hippocampus. Neuropharmacology. 2023;224.

35. Bluett RJ, Báldi R, Haymer A, Gaulden AD, Hartley ND, Parrish WP, et al. Endocannabinoid signalling modulates susceptibility to traumatic stress exposure. Nat Commun. 2017;8.

36. Huang GZ, Woolley CS. Estradiol acutely suppresses inhibition in the hippocampus through a sex-specific endocannabinoid and mGluR-dependent mechanism. Neuron. 2012;74:801–808.

37. Barker JM, Galea LAM. Sex and regional differences in estradiol content in the prefrontal cortex, amygdala and hippocampus of adult male and female rats. Gen Comp Endocrinol. 2009;164:77– 84.

38. Payrits M, Sághy É, Cseko K, Pohóczky K, Bölcskei K, Ernszt D, et al. Estradiol Sensitizes the Transient Receptor Potential Vanilloid 1 Receptor in Pain Responses. Endocrinology. 2017;158:3249–3258.

