## Supplemental Table 1 for "CB1R blockade unmasks TRPV1-mediated contextual fear generalization in female, but not male rats"

**Table S1: comprehensive statistics results**

| Figure | Test | Effect/comparison | Result | p value |
|---|---|---|---|---|
| 1B | Chi-square | Drug effect | Chi sq = 2.2 | P=0.53 |
| 1D | 2-way ANOVA | Trial x drug interaction | F (18, 276) = 1.110 | P=0.3414 |
| | | Main effect of trial | F (5.088, 234.0) = 0.7592 | P=0.5822 |
| | | Main effect of drug | F (3, 46) = 0.3702 | P=0.7749 |
| 1E | Mixed-effects model | Trial x drug interaction | F (18, 118) = 0.8232 | P=0.6700 |
| | | Main effect of trial | F (4.006, 78.79) = 2.573 | P=0.04* |
| | | Main effect of drug | F (3, 118) = 2.734 | P=0.047* |
| 1F | 2-way ANOVA | Trial x drug interaction | F (18, 324) = 0.7494 | P=0.7586 |
| | | Main effect of trial | F (4.953, 267.5) = 1.124 | P=0.3476 |
| | | Main effect of drug | F (3, 54) = 0.6473 | P=0.5880 |
| 2A (FC-BL) | 1-way ANOVA | Main effect of drug | F (3, 47) = 1.461 | P=0.2372 |
| 2A (FC-Tones) | 2-way ANOVA | Trial x drug interaction | F (18, 282) = 1.247 | P=0.2228 |
| | | Main effect of trial | F (5.022, 236.0) = 56.40 | P<0.0001 |
| | | Main effect of drug | F (3, 47) = 3.913 | P=0.0142 |
| | Dunnett's post-hoc | VEH vs. AM | | p range 0.003-0.03 trials 3*,4*,6*,7** all other comparisons ns. |
| 2A (EX-BL) | 1-way ANOVA | Main effect of drug | F (3, 47) = 7.461 | P=0.0003*** |
| | Dunnett's | AM vs. VEH | | P=0.0002*** |
| | Dunnett's | AM vs. CPZ | | P=0.0015** |
| | Dunnett's | AM vs. AM+CPZ | | P=0.0057** |

| | | | | |
|---|---|---|---|---|
| 2A (EX-Tone Blocks) | 2-way ANOVA | Trial x drug interaction | F (27, 423) = 1.062 | P=0.3834 |
| | | Main effect of trial | F (9, 423) = 14.05 | P<0.0001 |
| | | Main effect of drug | F (3, 47) = 4.018 | P=0.0126 |
| | Dunnett's | AM+CPZ vs. AM | | p range 0.0008-0.02 trials 3***,4**,5*,6* all other comparisons ns. |
| 2A (EXR-BL) | 1-way ANOVA | Main effect of drug | F (3, 47) = 5.437 | P=0.0027** |
| | Dunnett's | AM vs. VEH | | P=0.0077** |
| | Dunnett's | AM vs. CPZ | | P=0.0057** |
| | Dunnett's | AM vs. AM+CPZ | | P=0.0041** |
| 2A (EXR-Tone Blocks) | 2-way ANOVA | Trial x drug interaction | F (12, 188) = 0.8944 | P=0.5538 |
| | | Main effect of trial | F (3.342, 157.1) = 4.792 | P=0.0022** |
| | | Main effect of drug | F (3, 47) = 5.243 | P=0.0033** |
| | Dunnett's | AM vs. VEH | | P=0.02 on trials 1*,5* |
| | Dunnett's | AM vs. CPZ | | P=0.02 on trials 1*,5* |
| | Dunnett's | AM vs. AM+CPZ | | P range 0.004-0.03 on trials 1**, 2*, 3*, 4*, 5* |
| 2B (FC-BL) | 1-way ANOVA | Main effect of drug | F (3, 17) = 1.821 | P=0.1816 |
| 2B (FC-Tones) | 2-way ANOVA | Trial x drug interaction | F (18, 102) = 1.469 | P=0.1172 |
| | | Main effect of trial | F (3.965, 67.40) = 2.755 | P=0.0353* |
| | | Main effect of drug | F (3, 17) = 11.43 | P=0.0002*** |

| | | | | |
|---|---|---|---|---|
| | Dunnett's | AM+CPZ vs. VEH | | P range 0.004-0.01 on trials 2***, 4**, 5** |
| | Dunnett's | AM+CPZ vs. AM | | P range 0.0001-0.01 on trials 2****, 5** |
| | Dunnett's | AM+CPZ vs. CPZ | | P range 0.002-0.04 on trials 3*, 4**, 5**, 6* |
| 2B (EX-BL) | 1-way ANOVA | Main effect of drug | F (3, 17) = 3.413 | P=0.0414* |
| | Dunnett's | AM+CPZ vs. VEH | | P=0.02*<br>All other comparisons ns |
| 2B (EX-Tones) | 2-way ANOVA | Trial x drug interaction | F (27, 153) = 1.187 | P=0.2548 |
| | | Main effect of trial | F (3.914, 66.54) = 5.469 | P=0.0008*** |
| | | Main effect of drug | F (3, 17) = 1.690 | P=0.2070 |
| 2B (EXR-BL) | 1-way ANOVA | Main effect of drug | F (3, 18) = 2.378 | P=0.1037 |
| 2B (EXR-Tones) | 2-way ANOVA | Trial x drug interaction | F (12, 72) = 0.8340 | P=0.6156 |
| | | Main effect of trial | F (2.664, 47.96) = 1.925 | P=0.1443 |
| | | Main effect of drug | F (3, 18) = 0.3280 | P=0.8051 |
| 2C (FC-BL) | 1-way ANOVA | Main effect of drug | F (3, 54) = 0.9465 | P=0.4247 |
| 2C (FC-Tones) | 2-way ANOVA | Trial x drug interaction | F (18, 324) = 1.001 | P=0.4577 |
| | | Main effect of trial | F (5.093, 275.0) = 41.46 | P<0.0001**** |
| | | Main effect of drug | F (3, 54) = 3.000 | P=0.0384* |
| | Dunnett's | VEH vs. AM | | P=0.0065** trial 6 only |

| | | | | |
|---|---|---|---|---|
| | Dunnett's | VEH vs. AM+CPZ | | P=0.04 trial 6 only |
| 2C (EX-BL) | 1-way ANOVA | Main effect of drug | F (3, 54) = 1.124 | P=0.3477 |
| 2C (EX-Tones) | 2-way ANOVA | Trial x drug interaction | F (27, 486) = 1.046 | P=0.4038 |
| | | Main effect of trial | F (4.427, 239.1) = 18.56 | P<0.0001**** |
| | | Main effect of drug | F (3, 54) = 0.5748 | P=0.6340 |
| 2C (EXR-BL) | 1-way ANOVA | Main effect of drug | F (3, 54) = 1.930 | P=0.1357 |
| 2C (EXR-Tones) | 2-way ANOVA | Trial x drug interaction | F (12, 216) = 0.9654 | P=0.4830 |
| | | Main effect of trial | F (3.211, 173.4) = 3.831 | P=0.0093** |
| | | Main effect of drug | F (3, 54) = 0.8436 | P=0.4760 |
| 3B | Unpaired t-test | Effect of conditioning | t=2.602, df=18 | P=0.018* |
| | Unpaired t-test | Effect of conditioning | t=0.2164, df=18 | P=0.831 |
| | Unpaired t-test | Effect of conditioning | t=0.6415, df=18 | P=0.529 |
| 3C | Unpaired t-test | Effect of conditioning | t=0.5926, df=28 | P=0.558 |
| | Unpaired t-test | Effect of conditioning | t=1.563, df=26 | P=0.13 |
| | Unpaired t-test | Effect of conditioning | t=2.411, df=27 | P=0.023* |
| 4C | 2-way ANOVA | Sex x experience interaction | F (2, 41) = 0.9034 | P=0.4131 |
| | | Main effect of sex | F (1, 41) = 2.633 | P=0.1124 |
| | | Main effect of experience | F (2, 41) = 0.7036 | P=0.5007 |
| 4D | 2-way ANOVA | Sex x experience interaction | F (2, 41) = 0.9351 | P=0.4007 |
| | | Main effect of sex | F (1, 41) = 2.813 | P=0.1011 |
| | | Main effect of experience | F (2, 41) = 0.4887 | P=0.6170 |
| 4E | 2-way ANOVA | Sex x experience interaction | F (2, 41) = 0.5078 | P=0.6055 |
| | | Main effect of sex | F (1, 41) = 1.371 | P=0.2483 |

| | | | | |
|---|---|---|---|---|
| | | Main effect of experience | F (2, 41) = 0.2398 | P=0.7879 |
| 5H | 2-way ANOVA | Sex x labeling interaction | F (1, 24) = 0.1981 | P=0.6603 |
| | | Main effect of sex | F (1, 24) = 0.02454 | P=0.8768 |
| | | Main effect of labeling | F (1, 24) = 66.56 | P<0.0001**** |
| 5I | Welch's t-test | Main effect of sex | t=2.332, df=15.92 | 0.0332* |
